# Transform the Information Landscape Applying Waddington Landscape to Information Theory of Aging and Epigenetic Rejuvenation

**DOI:** 10.1101/2024.07.20.604423

**Authors:** Yue Wu

## Abstract

The Information Theory of Aging (ITOA) provides a framework for understanding aging and rejuvenation via changes in epigenetic information. These changes are visualized using Waddington landscape — a 3D topology that represents cell trajectories during changes in cell identity — constructed from field decomposition of scRNA velocity data. Studies have demonstrated that the Waddington landscape can serve as a mechanistic model of epigenetic information. It explains how age-driven decline in cellular function and reprogramming-induced rejuvenation can be understood by studying the transformations in the Waddington landscape. This study provides insights into the representation of epigenetic information and discusses how it can be reprogrammed to restore a younger cellular state.

## 1. Introduction

Almost all metazoans undergo aging—a process characterized by progressive functional decline and loss of homeostasis that ultimately leads to death. Studies have shown that transient expression of the reprogramming factors Sox2, Oct4, and Klf4 (OSK) can reset the cellular epigenetic landscape, reverse epigenetic aging, and rejuvenate metabolic function—an effect known as partial reprogramming-induced epigenetic rejuvenation (Ocampo et al., 2016; Browder et al., 2022; Lu et al., 2020). This highlights epigenetic rejuvenation as a promising research focus for potentially reversing aging. The Information Theory of Aging (ITOA) suggests that aging results from the loss of epigenetic information over time; however, the original information is retained (Lu et al., 2023; Yang et al., 2023b). OSK reprogramming can access this memory and restore youthful epigenetic states via epigenetic modifiers, leading to rejuvenation (Lu et al., 2023). This study explores ITOA using Waddington landscape, a 3D model that describes the travel trajectories of cells with changes in their cellular identities, to explain the aging process, as well as epigenetic rejuvenation.

## 2. Aging and Epigenetic Information

### 2.1. Life as information

Information refers to the extent to which a given informational relation alters or enhances existing knowledge (Marcos, 2011). The developmental and aging processes of an organism reflect a four-dimensional pattern of information that guides the dynamic evolution of phenotypes across three-dimensional space and time.

According to the central dogma of biology, DNA is transcribed into RNA, which is then translated into proteins that perform specific functions that shape phenotypes. Despite this, the DNA sequence of an organism remains almost unchanged throughout its lifespan. Epigenetics is the science that studies the molecules and mechanisms that maintain different gene activity states without altering DNA sequences (Cavalli and Heard, 2019). Epigenetic regulation includes DNA methylation, histone modification, chromatin remodeling, non-coding RNA (ncRNA) regulation, and RNA modification (Wang et al., 2022). These mechanisms control DNA expression dynamics and dictate the spatiotemporal order of life information.

Given the important role of epigenetics in the temporal development of phenotypes, it is not surprising that reprogramming-induced epigenetic rejuvenation is the only in vivo method capable of resetting both the transcriptome and the DNA methylome, facilitating long-term functional recovery (Lu et al., 2023). Currently, this is the only known method capable of achieving this resetting.

Before discussing the representation of such information, it is important to address the recurring question of where this information resides.

### 2.2. Location of epigenetic information

As reviewed by Cohen et al. (2022), biological aging exhibits several key features: (1) networks of interacting elements, including key metabolism pathways such as mTOR, Sirtuins, and AMPK (Bitto et al., 2015); (2) feedback/feedforward loops, exemplified by senescence-associated secretory phenotypes from paracrine signaling (Ren et al., 2009; Freund et al., 2010); (3) multi-scale or modular hierarchical structure, such as the propagation of dysfunctional aged cells across tissues (West, 2012; Kuo et al., 2020); (4) nonlinear dynamics, such as the accelerating decline in multiple systems at the end of life (Fried et al., 2021). These hallmarks illustrate that biological aging is a complex process (Cohen et al., 2022).

The act of digging down into the location of information is prone to losing sight of the entire complex system, starting at the organism level to determine where information resides in the tissue, narrowing to the tissue level to locate information within cells, further examining at the cellular level to explore molecular information, and ultimately studying atomic interactions. A characteristic of a complex system is that its emergent properties cannot be directly or additively inferred from its components (Cohen et al., 2022). Even if a clear source of aging information is found, the aged biological mechanics of the organism would be a deteriorating receiver to receive and process such information, which would not only undermine the causal link between the information source and aging phenotypes but also increase the entropy of the information source itself.

Thus, given the regulatory role of epigenetic information in the aging process illustrated in the previous section, a more direct and practical question to ask in aging theory is: how can we represent and manipulate epigenetic information?

## 3. Represent Epigenetic Information with Waddington Landscape

Waddington landscape describes the travel trajectories of cells with changes in their cellular identities. A classic example of this is stem cell differentiation. In this 3D model, a stem cell starts at a high point and rolls down, gradually losing its potential, and following specific pathways to differentiate into particular cell types (Waddington, 1957). More than just a metaphor, as originally proposed in 1957, the Waddington landscape has been shown to have a solid mathematical meaning through quasi-stability and vector field analyses of biological differentiation processes (Fusco et al., 2014; Bhattacharya et al., 2011).

Methods have been adopted to construct the Waddington landscape, such as transcriptome signaling entropy or Hopfield neural networks, to reflect the differential patterns of cells (Qin et al., 2023; Fard et al., 2016). Here, I used field decomposition of the uniform Manifold Approximation and Projection for Dimension Reduction (UMAP) embedding of RNA velocity vectors, incorporating epigenetic regulation of chromatin opening or closing status, to construct a Waddington landscape of 11,605 human hematopoietic progenitor stem cells (HPSCs) (**Fig.1**) (Jia and Chen, 2023; Li et al., 2023). Essentially, the greater the magnitude of the velocity vector in the dimensional decomposition vector fields, the steeper the landscape gradient along the z-axis. UMAP embedding of the RNA velocity vector field matches the known differentiation pathways of HPSCs (Li et al., 2023).

**Figure 1:**
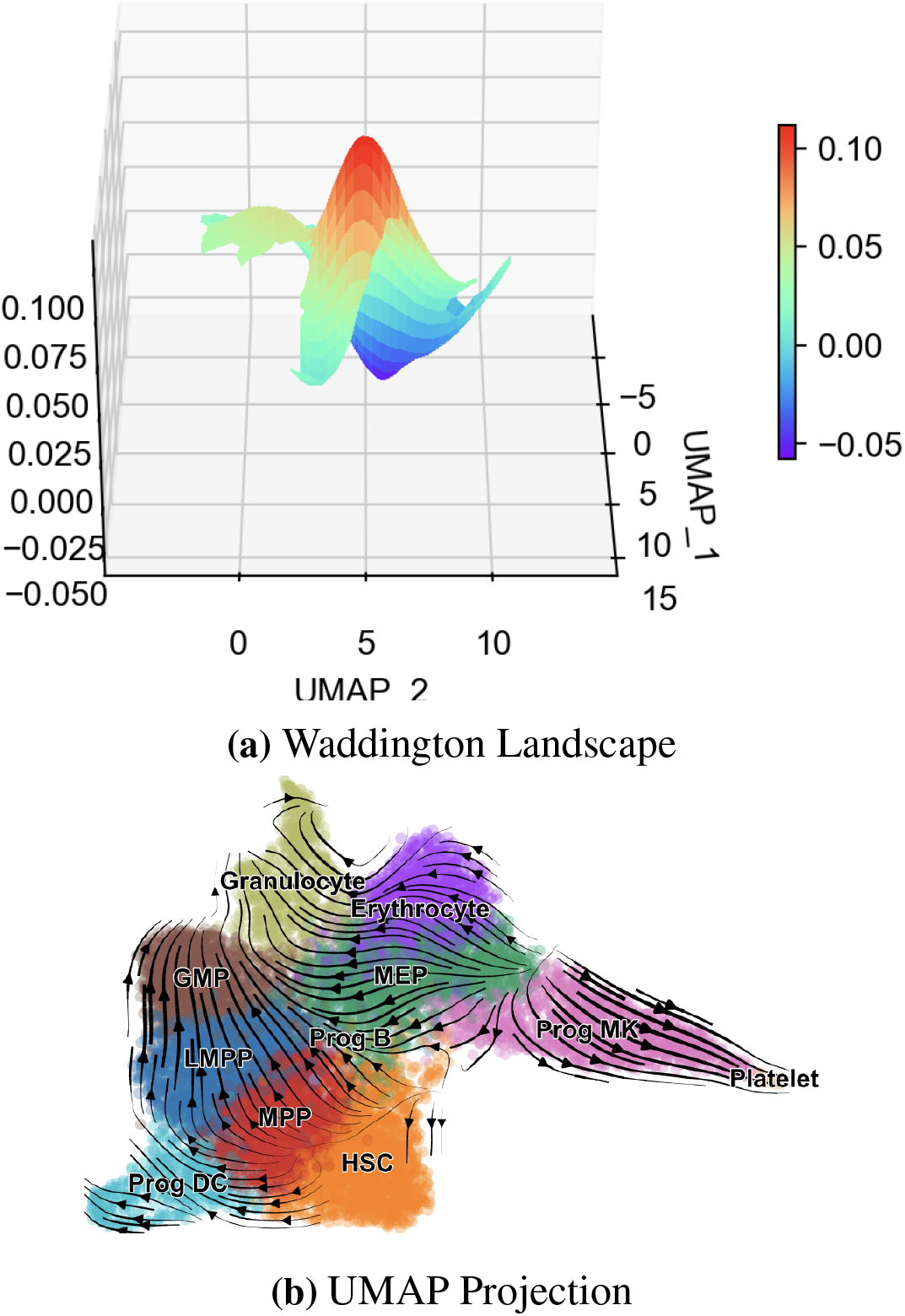
Computations of HSPC Differentiation from RNA Velocity Data

The top part of the landscape describes HPSC differentiation into erythroid and myeloid lineages, regulated by the transcription factors Pou1 and GATA5, as described in a mathematical model (Huang et al., 2007). The corresponding portion of the UMAP integrity vector field fits this mathematical model with parameters within a meaningful range, and the landscape reflects the differentiation trajectories of bidirectional attractors from an unstable equilibrium point (**Fig.2**) (Huang et al., 2007). This illustrates that the cellular differentiation process is separated into two attractor sites corresponding to distinct phenotypes (erythroid and myeloid lineages). This demonstrates that the Waddington landscape can serve as a mechanistic model for the different stages of an organism’s development.

**Figure 2:**
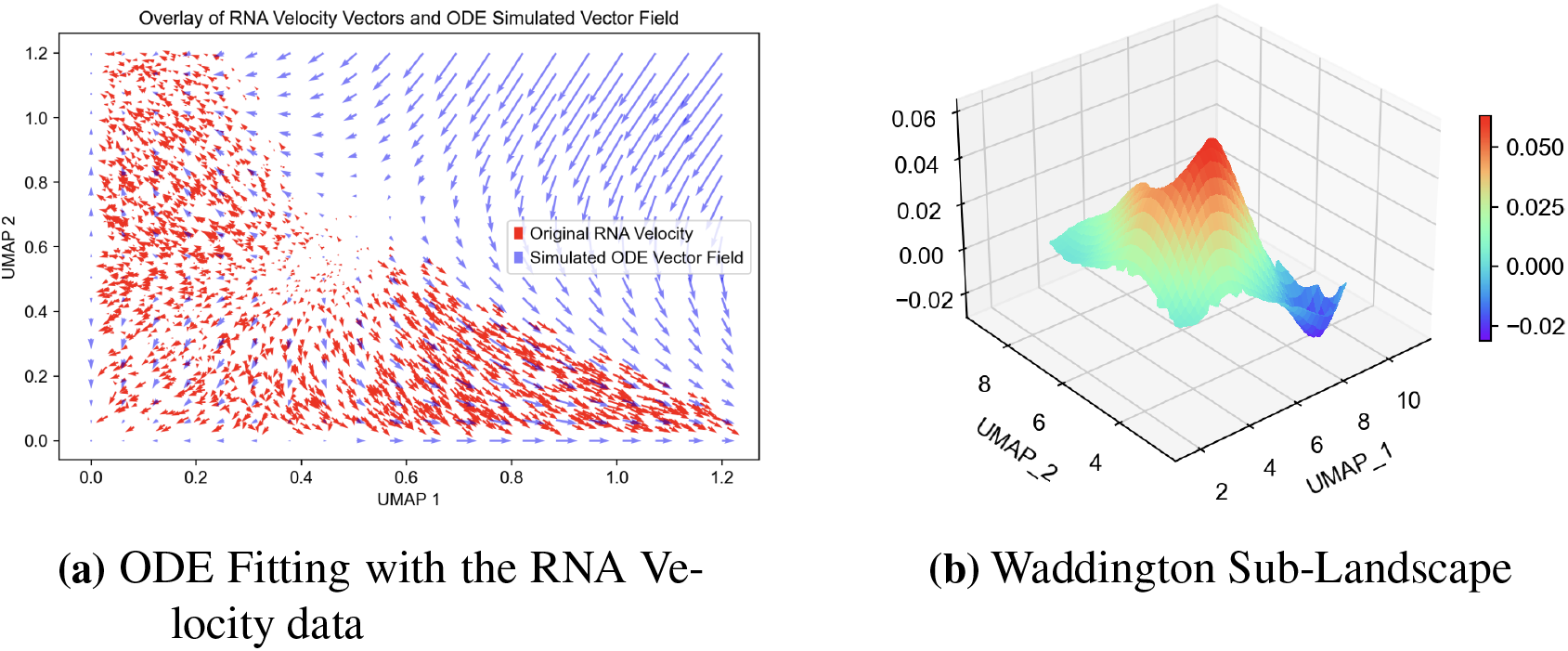
Computations of Myeloid and Ethyroid Lineage Differentiation

The Waddington landscape can be extended to include the aging stage of differentiation, which refers to the process by which aged cells lose their somatic identity and cannot perform their correct functions (Ono and Cutler, 1978; Cutler, 1982; Lu et al., 2023). Reflecting the landscape is the erosion of the attractor topology that originally held the cells in the correct somatic landscape, and the cells traveled down to a region with an ambiguous identity (**Fig.3**). The proposed mechanism is that DNA double-strain breaks, viral infections, and physical and other damages induce the relocalization of chromatin-modifying proteins (RCM) to address these issues (Yang et al., 2023b; Oberdoerffer et al., 2008; Poganik et al., 2023; Lu et al., 2020; Karg et al., 2023). After RCM, the chromatin-modifying protein positions remain without returning to the original states, thereby introducing noise to the original epigenetic information, leading to the blurring of the landscape (Oberdoerffer et al., 2008; Lu et al., 2023). This aligns with the evolutionary aging theory of antagonistic pleiotropy, which suggests that epigenetic noise benefits cells in the short term by recruiting chromatin factors to repair DNA damage and keep the cells alive. However, in the long term, these actions are detrimental and drive aging owing to the loss of epigenetic information (Williams, 1957; Lu et al., 2023; Yang et al., 2023b). Thus, the Waddington landscape can represent epigenetic information given its alignment with the mechanisms and phenotypes of biological aging.

**Figure 3:**
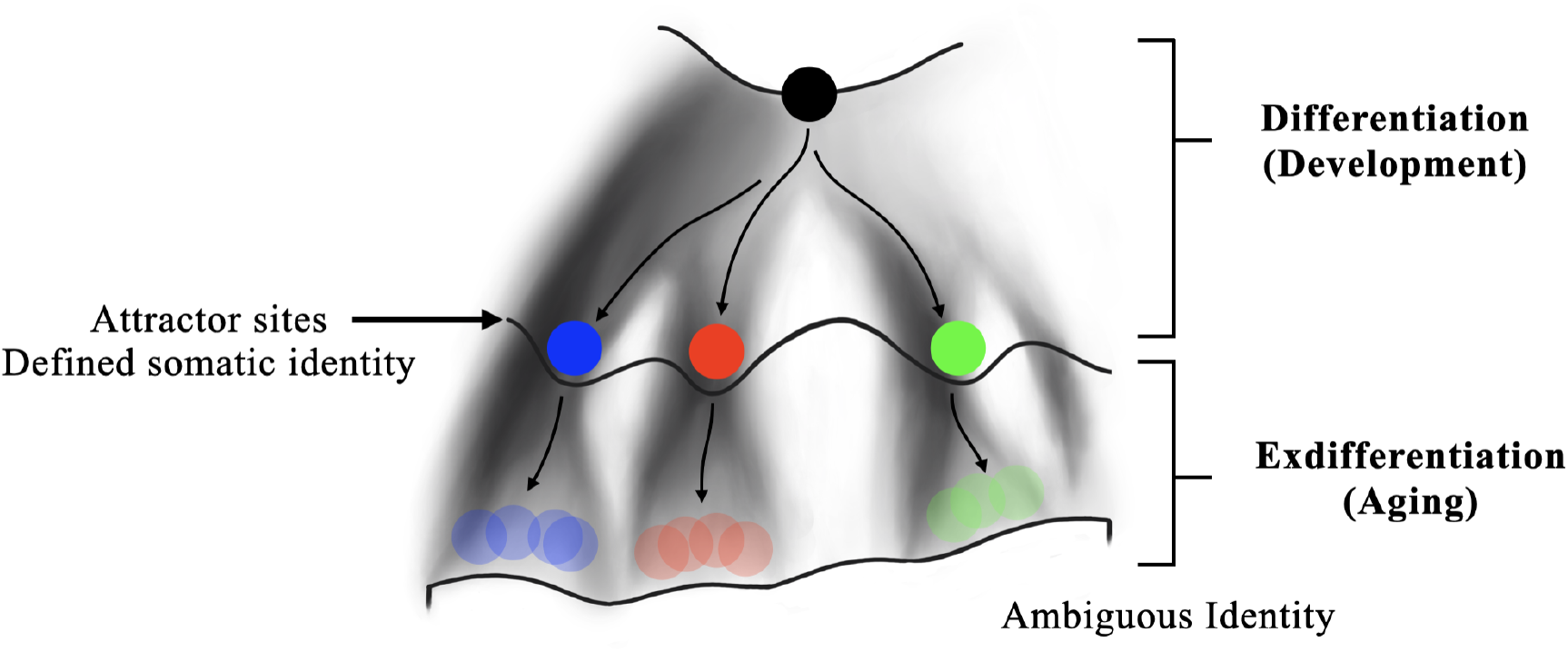
Complete Waddington Landscape of Development and Aging

## 4. Transformation of Epigenetic Information Landscape

### 4.1. The reprogramming process assesses the memory of the epigenetic landscape and transforms it

Without correcting mutations in the DNA sequence, the biological age resets in zygote formation and the normal lifespans of cloned mammals from somatic nuclear transfer suggests that aged cells possess the memory of youthful epigenetic information (Gladyshev, 2021). As previously mentioned, such memories encompass a complex system of biological aging.

Recent studies have suggested that partial reprogramming induced by Sox2, Oct4, and Klf4 (OSK) rewrites the epigenetic landscape in vivo and improves the cellular metabolic performance (Ocampo et al., 2016; Browder et al., 2022; Gill et al., 2022; Lu et al., 2020; Sarkar et al., 2020). In other words, the OSK acquires access to epigenetic information and transforms the landscape into rejuvenated cells.

Camillo and Quinlan proposed that during partial reprogramming, the Waddington landscape is flipped and flattened (De Lima Camillo and Quinlan, 2021). The number of stem cells was found to be maximum in the untransformed landscape. However, during reprogramming, the cells move toward the stem cell state following the reverse landscape gradient, flipping the maxima into minima, and vice versa (**Fig.4**). Taking a mathematical perspective on the computed landscape, the RNA velocity vector reversed its direction from moving away from the progenitors to moving toward the progenitors. Thus, the landscape derived from vector field decomposition will also be inverted (Jia and Chen, 2023). Additionally, the observation that reprogramming generates cancer cells with a much higher eAge than the organism and iPSC with a much lower eAge implies that the landscape is flattened (Horvath, 2013). When OSK is applied to cells, it calls out the memory of epigenetic information and transforms the epigenetic landscape by reversing or flipping it.

**Figure 4:**
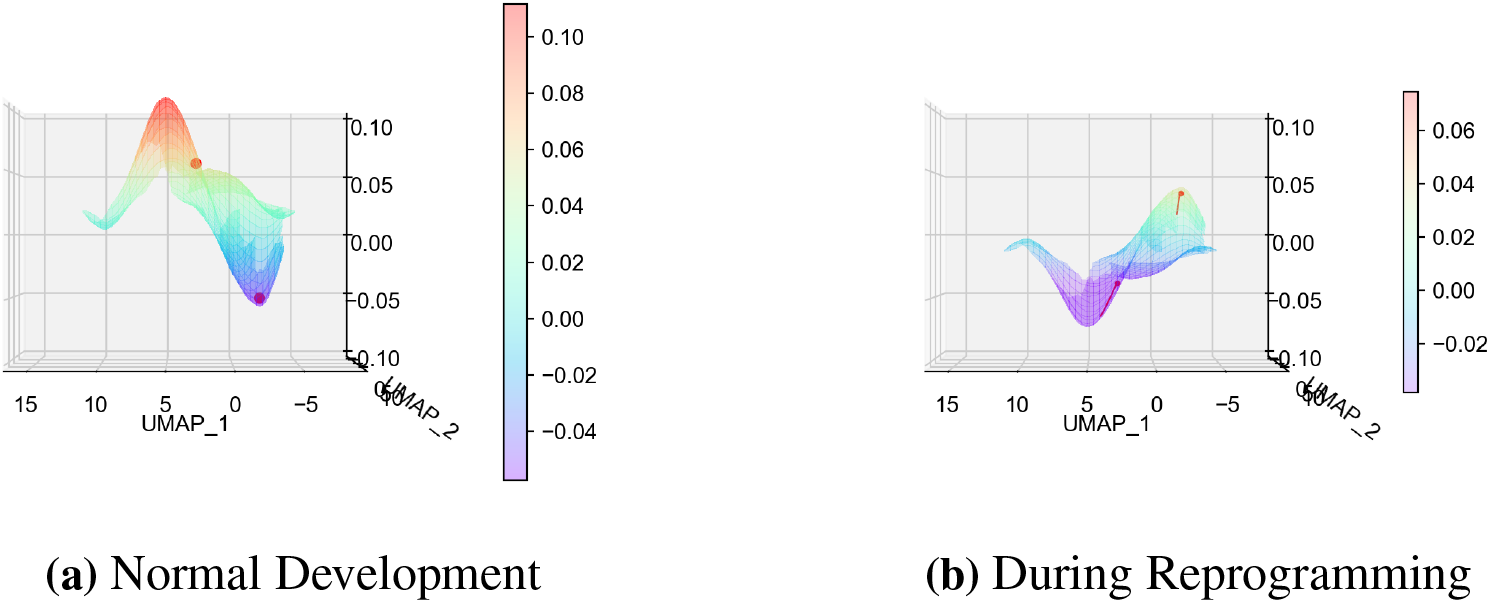
The reverse and flattening of HSPC differentiation Waddington landscape. The red dotes represent the epigenetic status of two cells. (b) shows the trajectories of cells traversing through the landscape.

Thus, aged ex-differentiated cells would travel down to the younger attractor site with a defined somatic identity. Cells that are already in the attractor site, if given enough time and conditions, would travel out of the attractor influence toward their progenitor state and transiently enter a zone of epigenetic instability and ambiguous identity (Sarkar et al., 2020). As reprogramming ended, the Waddington landscape was restored. The previously old cells remained at their position due to the influence of the attractor, and the young cells slid back to the attractor stage (**Fig.5**).

**Figure 5:**
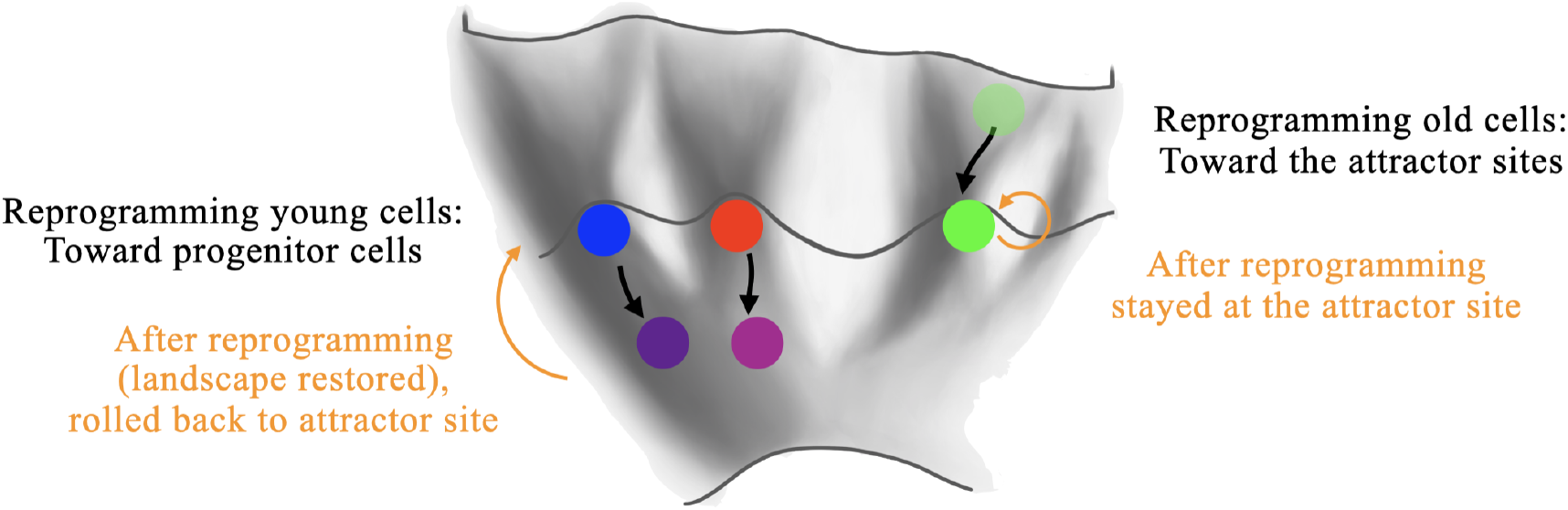
Waddington Landscape during Reprogramming

Their flattened nature allows cells to surpass barriers more easily and traverse faster, which makes reprogramming an effective and sensitive method to move cells across the epigenetic landscape, as illustrated by experiments with mature lung epithelial cells, where partial reprogramming created an unnatural progenitor state (Guo et al., 2017). This shows that the transformation of epigenetic information can cause cells to settle into different local minima that do not align with their original differentiation trajectory, thereby enabling transdifferentiation.

### 4.2. The access and transformation of epigenetic information by non-reprogramming biological features

Experimental observations validate factors or biological features, in addition to reprogramming factors, to gain access to and the capacity to transform epigenetic information in different ways.

1. **Reinforcing attractor landscape to enhance cellular identity preservation**. For example, active transcription-associated chromatin marks such as DOT1L-catalyzed H3K79 methylation and FACT-mediated histone turnover act to maintain somatic gene expression profile while inhibiting reprogramming (Arabacı et al., 2021). Another example is metformin, an activator of the longevity pathway AMPL, which increases lifespan in mice (Martin-Montalvo et al., 2013). 100 M concentration of metformin is sufficient to alleviate several aging markers in human cells, but only 10 M significantly reduces reprogramming efficiency (Kajbaf et al., 2016; Fang et al., 2018; Vazquez-Martin et al., 2012). As the treatment fosters genetic stability by deepening the grooves in the Waddington landscape, when the landscape is flipped during reprogramming, these deeper grooves become higher barriers for cells to cross (De Lima Camillo and Quinlan, 2021). This explains why some longitudinally promoted treatments oppose reprogramming-induced rejuvenation.
2. **Carving new horizontal paths**. Reprogramming research in *C. elegans* has achieved direct transdifferentiation in vivo, in which one tissue is directly converted into another from a different germ layer lineage (Riddle et al., 2016). The utilized transcription factors create a new path in the landscape, which leads the cells to traverse without traveling toward their corresponding progenitors.
3. **Carving new vertical paths**. Sodium butyrate and alpha-ketoglutarate have been shown to increase reprogramming efficiency while preserving cell identity (Yang et al., 2023a). Also, an alternative reprogramming method, 7F reprogramming with seven factors Jdp2–Jhdm1b–Mkk6–Glis1–Nanog–Essrb–Sall4, activates another reprogramming pathway different from the OSK-method, and is suggested to have potential in decoupling rejuvenation and dedifferentiation (Wang et al., 2019; Yu cel and Gladyshev, 2024). These factors may carve a straightforward path in the reprogramming landscape to rejuvenate cells without the risk of acquiring an undefined identity. Identifying such factors is an important target for rejuvenation studies to mitigate the safety risk of cellular identity obfuscation.
4. **Formation of new attractor sites**. Mutations in oncogenic DNA sequences can create new attractor sites at unstable locations within the landscape. For example, during the differentiation of megakaryocyte erythroid progenitors (MEP) into megakaryocyte cell types, cells travel down from local maxima to attractor sites of defined cell identity, following the potential gradient of the epigenetic landscape. However, mutations in oncogenic DNA sequences alter the landscape and create new attractor sites (corresponding to leukemia) on the potential gradient between MEP and defined blood cell types. This causes leukemia cells to self-replicate stably with less functional cellular identity (Jordan, 2004). Consider acknowledging the limitations of this study before the conclusion section.

## 5. Conclusion

This study illustrates that the Waddington landscape can mechanistically represent epigenetic information that instructs the dynamics of development and aging processes. OSK reprogramming factors achieve rejuvenation by accessing the memories of epigenetic information, reversing and flattening the landscape, and allowing cells to travel to a younger state. Other biological entities can also transform the landscape in various ways to achieve different effects, such as stabilizing cell identity, transdifferentiation, and cancer formation.

Although this study mainly described aging mechanisms at the cellular level, developing effective aging interventions requires an understanding of the information theory of aging from a comprehensive, systematic perspective. At the tissue level, it has been found that adult mouse organs are composed of mosaics of cells spanning various ages (Arrojo E Drigo et al., 2019). Exploring how the heterogeneity of epigenetic information in tissue cells contributes to the entropy of tissue-level epigenetic information and how this entropy influences tissue aging is an interesting avenue for further research. At the organismal level, intra-organ positive feedback and feed-forward loops sustain and stabilize phenotypic states, and circulating factors in the blood can induce body-wide phenotypic changes (Kang and Yang, 2020). Instead of focusing on a single molecular pathway, examining network interactions and representing epigenetic information from a comprehensive and systematic perspective could be key research focuses in the study of aging.

## Notes

### Competing Interest Statement

The authors have declared no competing interest.

### Summary of Updates

Language is revised to a more academic style

